# Engineering efficient termination of bacteriophage T7 RNA polymerase transcription

**DOI:** 10.1101/2021.12.09.471997

**Authors:** Diana G. Calvopina-Chavez, Mikaela A. Gardner, Joel S. Griffitts

## Abstract

The bacteriophage T7 expression system is one of the most prominent transcription systems used in biotechnology and molecular-level research. However, T7 RNA polymerase is prone to read-through transcription due to its high processivity. As a consequence, enforcing efficient transcriptional termination is difficult. The termination hairpin found natively in the T7 genome is adapted to be inefficient, exhibiting 62% termination efficiency in vivo and even lower efficiency in vitro. In this study, we engineered a series of sequences that outperform the efficiency of the native terminator hairpin. By embedding a previously discovered 8-nucleotide T7 polymerase pause sequence within a synthetic hairpin sequence, we observed in vivo termination efficiency of 91%; by joining two short sequences into a tandem 2-hairpin structure, termination efficiency was increased to 98% in vivo and 91% in vitro. This study also tests the ability of these engineered sequences to terminate transcription of the *Escherichia coli* RNA polymerase. Two out of three of the most successful T7 polymerase terminators also facilitated termination of the bacterial polymerase with around 99% efficiency.

## INTRODUCTION

Protein expression systems derived from bacteriophage T7 are commonly used for high-level expression of recombinant proteins in engineered strains of *Escherichia coli* (Conrad et al. 1996; Lebedeva et al. 1994; Tabor 2001; Wycuff and Matthews 2000). In these systems, a protein-coding gene of interest is placed downstream of an 18 bp promoter that is specifically recognized by T7 RNA polymerase (T7RNAP) (Rong et al. 1998). This construct is then introduced into an *E. coli* strain that has been modified to carry a copy of the T7RNAP gene on its chromosome. Expression of T7RNAP is typically controlled by a chemically inducible bacterial promoter, such as the *lac* promoter (Lebedeva et al. 1994; Rong et al. 1998). To enhance chemical control of the system, a *lac* operator may also be installed downstream of the T7 promoter on the expression plasmid (Dubendorf and Studier 1991; Studier 1991; Studier and Moffatt 1986). T7RNAP transcribes DNA at a higher rate than *E. coli* RNA polymerase (EcRNAP) (Golomb and Chamberlin 1974; Iost et al. 1992), but its high processivity can result in read-through transcription and even circumnavigation of the entire expression plasmid. T7RNAP-generated transcripts are therefore often much longer than necessary, resulting in overexpression of unwanted proteins (McAllister et al. 1981; Tabor and Richardson 1985).

The native T7 terminator (T7nat) is found in the T7 phage genome between genes 10 and 11 (Dunn and Studier 1983), and its sequence and predicted structure are given in Table 1 and Figure S1, respectively. This terminator features a 15-bp stem with 5 G-U base pairs, a 6-nt loop, and a 3’ poly-U tract. T7nat is tuned to be inefficient in its native context, allowing upstream genes to be expressed at higher levels than the more modestly expressed downstream genes (Dunn and Studier 1983). Read-through transcription is thus built into the T7 genetic program to the benefit of the organism. In most protein expression plasmids, T7nat is found within a larger sequence context that promotes stronger termination efficiency than is accommodated by the core hairpin sequence described above (Kwon and Kang 1999; Song and Kang 2001). In one study, termination efficiency was found to decrease by half when the 100-bp sequence upstream of the terminator hairpin was omitted (Yoo and Kang 1996). This has prompted a search for more compact sequences facilitating stronger T7RNAP termination. In one such study, a modified T7 terminator sequence employing a more structurally favorable UUCG loop, as well as replacement of certain G-U base pairs with G-C base pairs, yielded a 40% improvement in termination efficiency in vitro (Mairhofer et al. 2015).

**Table 1.**
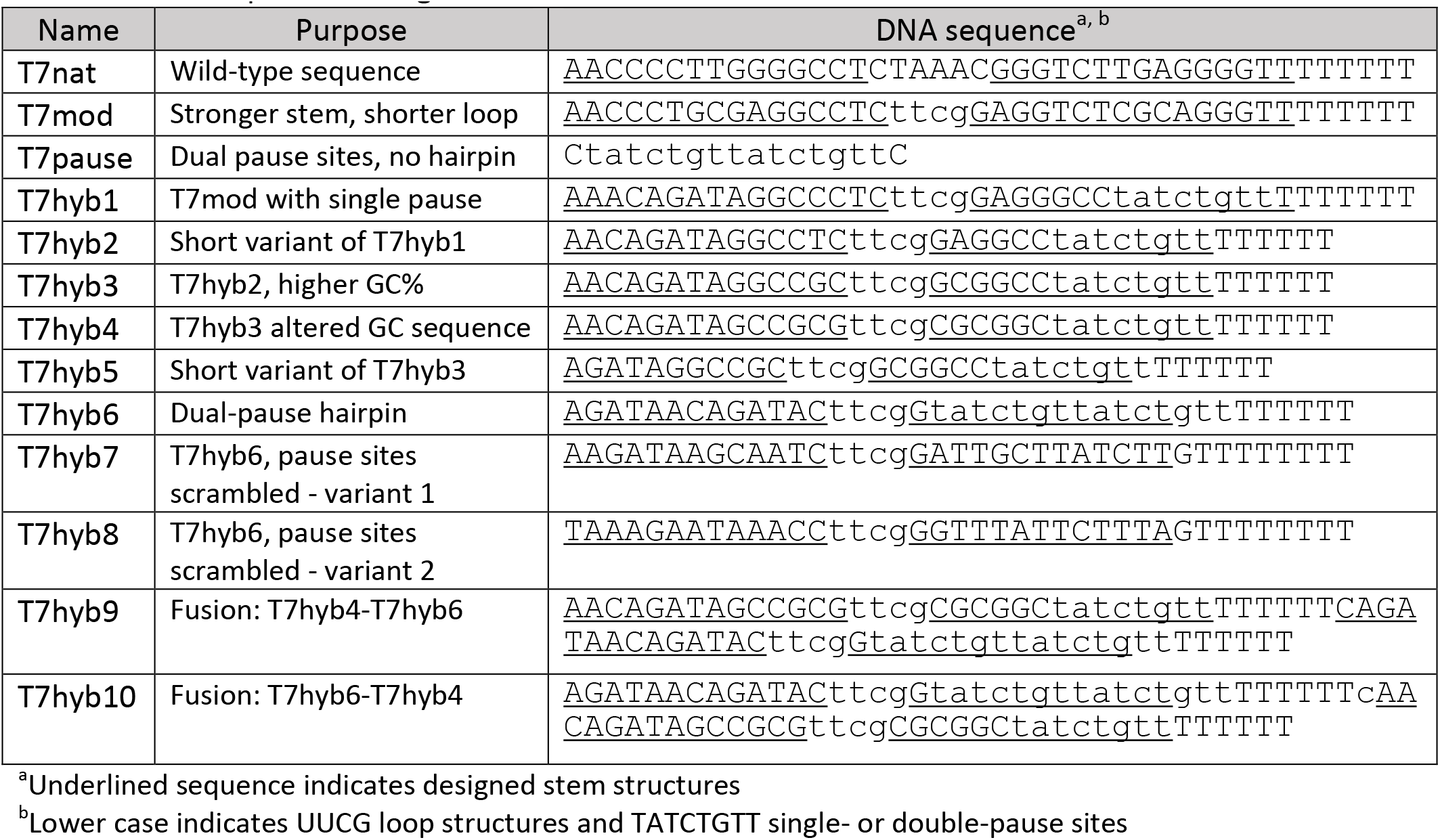
DNA sequence of engineered terminators.

In addition to hairpin-forming (Class I) T7RNAP terminators, unstructured and sequence-dependent (Class II) termination signals have been identified (Du et al. 2012; Macdonald et al. 1994), primarily through in vitro assays on non-phage DNA templates. One such sequence (TATCTGTT), stimulates T7RNAP pausing, which potentiates termination under certain circumstances (Barr and Wertz 2001; Lyakhov et al. 1998; Whelan et al. 1995). In this report, we show how hybrid sequences containing overlapping Class I and Class II elements give rise to compact and efficient synthetic T7RNAP termination signals.

## MATERIALS AND METHODS

### Bacterial strains, plasmids, and media conditions

All plasmids and strains used in this study are summarized in Table S1 and S2. For cloning purposes, all plasmids were first transformed into DH5a *E. coli* cells. Plasmids were then extracted and transformed into MG1655 WT *E. coli* cells for measuring transcription termination by fluorescence (see below). Cultures were grown at 37°C in Luria broth medium supplemented with chloramphenicol (30 mg/ml) and ampicillin (100 ug/ml) or only ampicillin (100 ug/ml) as appropriate.

### Plasmid construction for each terminator

A unique pair of oligonucleotides was hybridized to create the inserts containing each terminator sequence (Table S3). Hybridized oligos were designed to have sticky ends complementary to *Bam*HI and *Kpn*I. Oligo sequences are detailed in Table S3. For oligo hybridization, TEN buffer (5 mM Tris pH8.0, 0.5 mM EDTA pH8.0, 50 mM NaCl) was prepared and top and bottom oligos were added to a final concentration of 10uM each. The mixture was then brought to: 95°C for 1 min, 90°C for 15 min, 75°C for 20 min, 70°C for 20 min, 65°C for 20 min, 60°C for 20 min, 55°C for 20 min, and then held at 4°C. Parent plasmid pJG1113 containing GFP-*Bam*HI-*Kpn*I-RFP was digested with *Bam*HI and *Kpn*I, and inserts with each terminator were ligated.

### RNAP transcriptional termination measurements in vivo

Plasmids were transformed into *E. coli* MG1655 and reporter gene expression was measured after culturing strains for 8 hours at 37°C. To measure T7 RNAP termination, basal level expression of T7RNAP (with no arabinose added) was relied upon in these experiments. OD_600_ measurements were taken to adjust cell density in each culture prior to fluorescence measurements. After subtracting background fluorescence (based on a control strain lacking T7RNAP), the red/green ratio from the no-terminator control strain allowed for normalization to 1. This was compared to all red/green ratios from the terminator constructs to determine percent termination efficiency. For measurements of *E. coli* RNAP termination, similar conditions were used, except cultures were supplemented with 0.3 mM IPTG.

### In vitro transcription reactions

Linear terminator containing DNA templates were amplified from corresponding plasmids (see table S1). The forward primer (oDC81) had the T7 promoter sequence appended, and it hybridized around 100 bp upstream of the terminator; the reverse primer (oDC82) was designed to hybridize around 100 bp downstream of the terminator. After DNA amplification and purification, transcription reactions were carried out in a volume of 20 μl of RNase free water, 1x RNase buffer, 0.5 mM NTPs, 1 U/μl RNase inhibitor, 1 μg of linear DNA template, 2U/μl T7RNAP (NEB M0251S). Reactions were incubated at 37°C for 3 h. They were then placed at 75°C for 10 min to inactivate the enzyme. Reactions were treated with DNAse I (1U/μl) at 37°C for 20 min. RNA products were concentrated using the RNA Clean & Concentrator kit from Zymo (R1015), and 15 μl of sample was resolved in an 8% polyacrylamide gel in the presence of 7M Urea and 3ug/ml ethidium bromide (Figure S2). RNA products were visualized in a UV gel imager. Termination efficiency percentages were obtained by calculating the ratio of the integrated density of the terminated (lower molecular weight) band over the integrated density of the entire lane. Integrated densities were obtained with ImageJ software.

## RESULTS AND DISCUSSION

### Assay for measuring T7RNAP termination in vivo

A two-plasmid system was used to monitor T7RNAP termination efficiency in vivo, where plasmid pJG1115 expresses the gene for T7RNAP using the L-arabinose-inducible (ParaBAD) promoter. The second plasmid (pJG1113) contains a bicistronic cassette encoding a green fluorescent protein (msfGFP; hereafter referred to as GFP) and a red fluorescent protein (mScarlet-I; hereafter referred to as RFP) in which the restriction sites *Bam*HI and *Kpn*I are used to ligate terminator candidates between the two fluorescent protein reporters. This biscistronic cassette is expressed using the T7 promoter (P_T7_) (see Figure 1A for complete plasmid maps; see Supplemental Material file for complete DNA sequence of parent plasmids used in this study). GFP/RFP ratiometric analysis allows T7RNAP termination efficiency to be calculated, using control strains with no T7RNAP and with no terminator sequence (see materials and methods).

**Figure 1.**
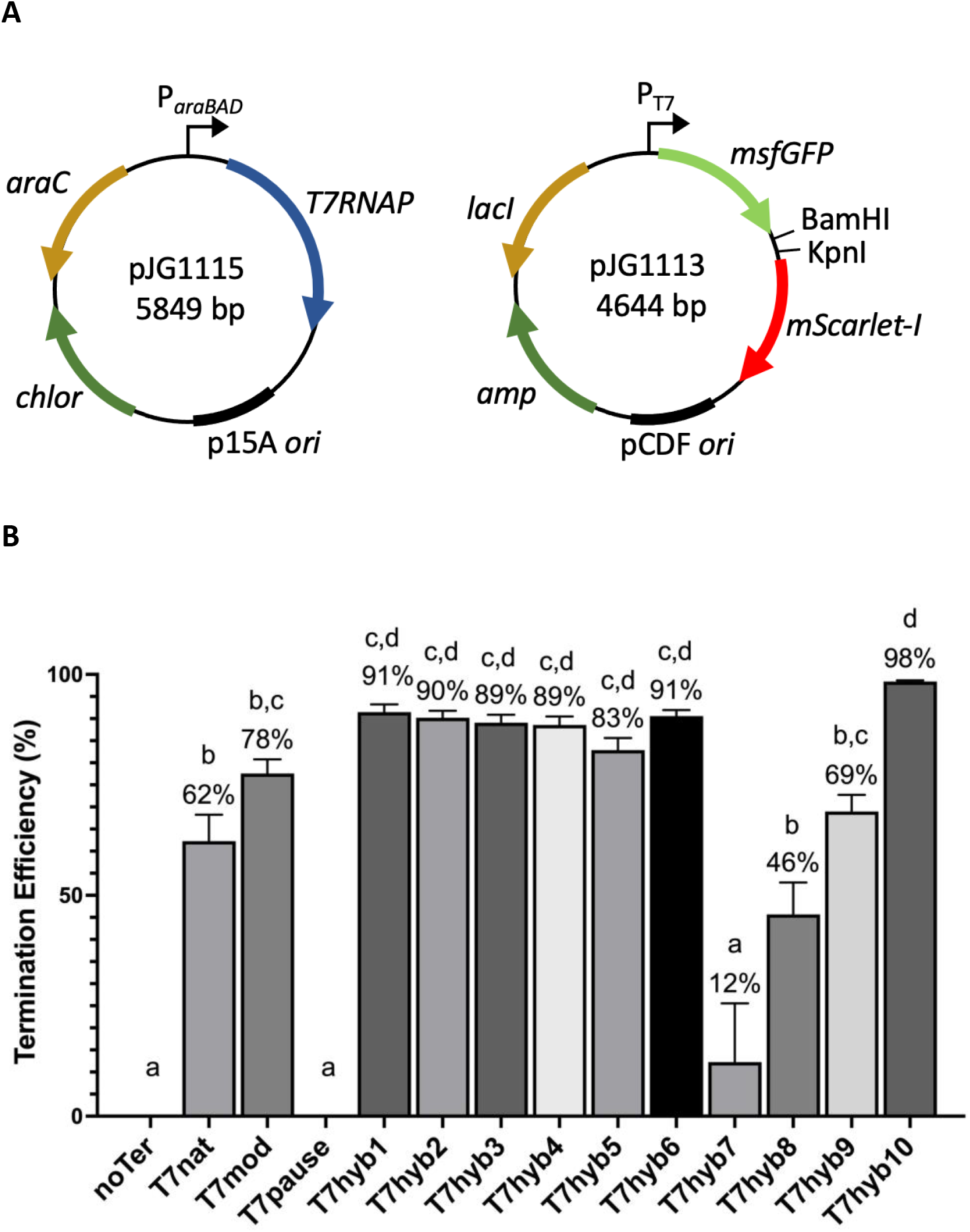
In vivo termination efficiency of designed T7 transcriptional terminators. (A) A two-plasmid system allows T7RNAP expression from an arabinose-inducible promoter and monitoring of T7RNAP-dependent transcription from a T7 promoter driving green and red fluorescent proteins. (B) Error bars represent standard deviation from the mean. Different letters denote statistically significant differences (P < 0.05) according to a Tukey multiple comparison test.

### Design of synthetic terminators

T7nat was used as a reference sequence throughout this study. A potentially more effective variant of T7nat, termed T7mod, was inspired by the UUCG loop-containing sequence described above (Mairhofer et al. 2015). These are both Class I terminators. As a Class II terminator sequence, we employed a known pause site (T7pause), with the sequence TATCTGTT, and this was inserted as two copies that overlap by one base pair (see Table 1). In remaining test sequences (T7hybX, where X is 1-10), Class I and Class II elements were combined, with pause sequences embedded in the poly-U-proximal segment of the terminator stem. T7hyb1-5 contain a single pause site and differ in stem length and stem %GC; T7hyb6 contains two slightly overlapping pause sites; T7hyb7 and T7hyb8 mimic T7hyb6, forming perfect hairpins but with the double-pause sequences being scrambled such that overall base composition was not altered. Tandem double-hairpins were also tested (T7hyb9 and T7hyb10). These designs allowed us to evaluate the influence of several structural and sequence parameters, including the embedded pause sequences. These sequences are summarized in Table 1, and Table S3 details the synthetic oligonucleotides used to clone these sequences.

### In vivo and in vitro performance of terminator designs

T7nat, T7mod, T7pause, and T7hyb1-10 were ligated into the GFP-RFP reporter plasmid and assayed for in vivo termination efficiency (see Table S1 and S2 for plasmids and strains details). The results of these tests are given in Figure 1B. T7nat exhibits 62% termination efficiency, with T7mod having modestly increased activity. T7pause shows no detectable activity in this assay. This is not surprising as previous reports have shown that Class II pause sites facilitate termination much more efficiently in vitro than in vivo (Du et al. 2009; Du et al. 2012). Where a single pause site is incorporated in the hairpin (T7hyb1-5), efficiency varies from 83% to 91%, with the short-stem variant (T7hyb5) showing the lowest efficiency. The double-pause design (T7hyb6) provides 91% efficiency despite the much lower GC content in the stem. To test whether the TATCTGTT pause sequences in T7hyb6 are critical, they were scrambled, resulting in molecules that would fold similarly, with identical %GC to T7hyb6. These scrambled variants (T7hyb7 and T7hyb8) exhibited much lower termination efficiency (12% and 46%, respectively), indicating that nucleotide order in these pause sequences strongly contributes to the efficiency of T7hyb6, even though the double-pause sequence on its own (T7pause) is ineffective. Finally, tandem double-hairpins were tested. The efficiency of T7hyb9 (tandem T7hyb4-T7hyb6) was lower than either single hairpin alone, while T7hyb10 (tandem T7hyb6-T7hyb4) was the most efficient of the whole set, at 98%. Why two single-hairpin elements joined in two possible orientations leads to such different termination efficiencies is unclear.

In vitro tests were carried out on a subset of the terminator sequences using recombinant T7RNAP (New England Biolabs) and PCR-generated DNA templates. Transcription products (size range: 127-312 nt) were resolved on polyacrylamide-urea gels and stained with ethidium bromide. Ratiometric analysis of terminated and run-off products was performed using Image J (see Table S4, S5 and S6 for integrated density values obtained from ImageJ for ratiometric analysis). In these tests, the relative efficiencies of terminator sequences reflected those determined in vivo, though calculated in vitro values were considerably lower (see Figure 2). Notably, however, the T7pause sequence did register a detectable amount of termination, and T7hyb6 (with its double pause) showed significantly enhanced termination over T7hyb1 (with only a single pause), while T7hyb1 and T7hyb6 were equivalently effective in vivo. We interpret this to mean that the pause sites, either alone or embedded in a hairpin, contribute more to termination efficiency in vitro than in vivo. This interpretation is supported by previous work (Du et al. 2012). As in the in vivo analysis, T7hyb10 was the best-performing design.

**Figure 2.**
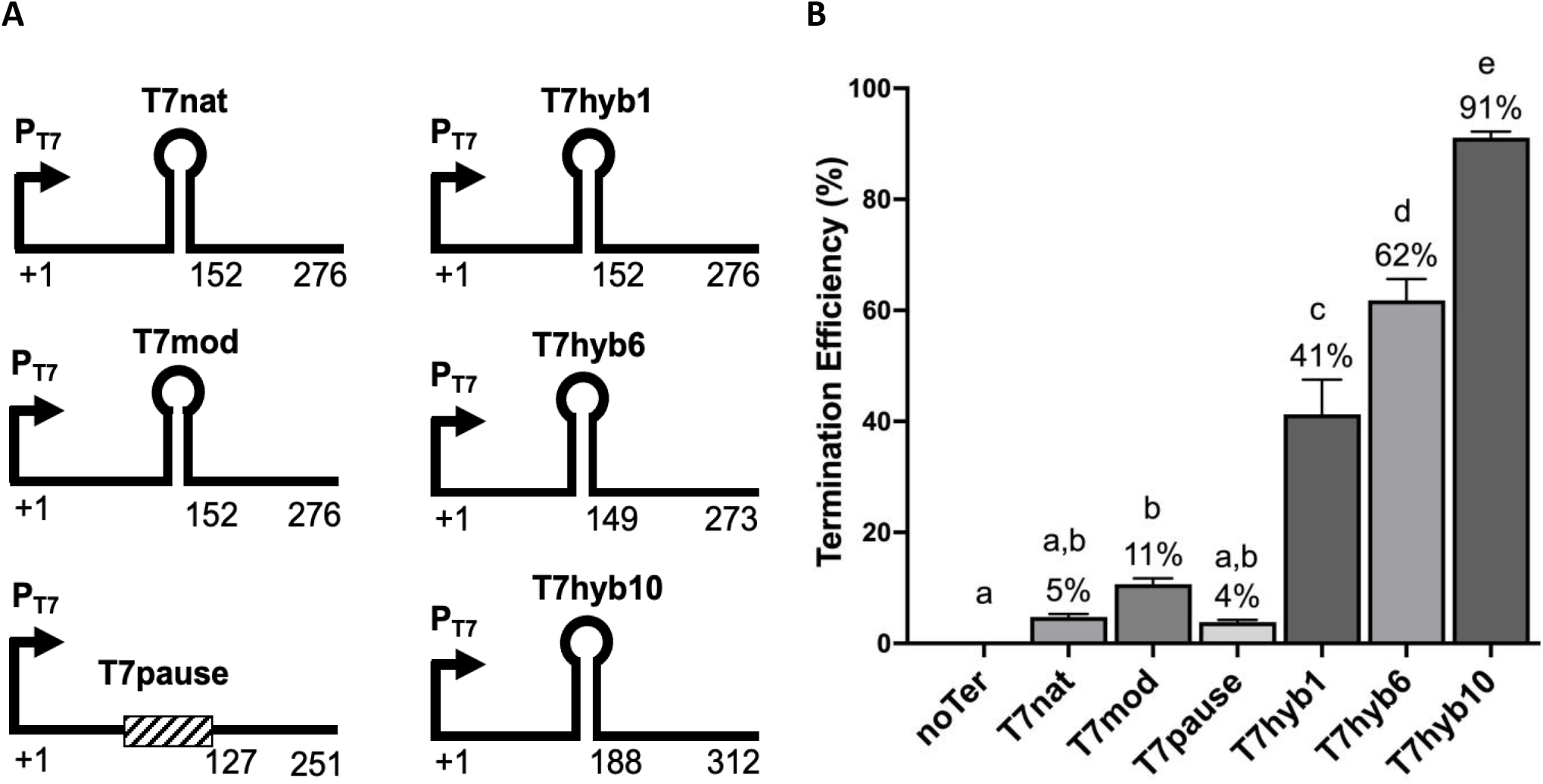
Performance of selected terminators in vitro. (A) Diagram of transcription templates for in vitro testing using recombinant T7RNAP. The three numbers underneath each diagram show the transcription initiation site, the predicted site of termination, and the length of the run-off product. Hairpin symbols represent the cloned T7 terminators. (B) In vitro termination efficiency of selected terminators. The error bars represent standard deviation from the mean. Different letters denote statistically significant differences (P < 0.05) according to a Tukey multiple comparison test.

### Terminator activity for the *E. coli* bacterial RNAP

To determine whether these synthetic terminators would be more broadly applicable in *E. coli*-based expression systems, they were tested in the context of the native bacterial polymerase (EcRNAP). To do this, pJG1113 (see Figure 1A) was modified to use the *lac* promoter to drive the GFP-RFP cassette (pJG1118; see Figure 3A). The same terminator sub-set used in the in vitro analysis was investigated here. As shown in Figure 3B, T7hyb1 and T7hyb10 gave outstanding termination efficiencies (99%) for EcRNAP in vivo. For all the data taken together, the tandem hairpin T7hyb10 sequence is an excellent all-around termination signal for RNA polymerases from both T7 (98%) and *E. coli* (99%). For a simpler single-hairpin alternative, T7hyb1 performs best, with in vivo efficiency values of 91% for the phage polymerase and 99% for the bacterial polymerase.

**Figure 3.**
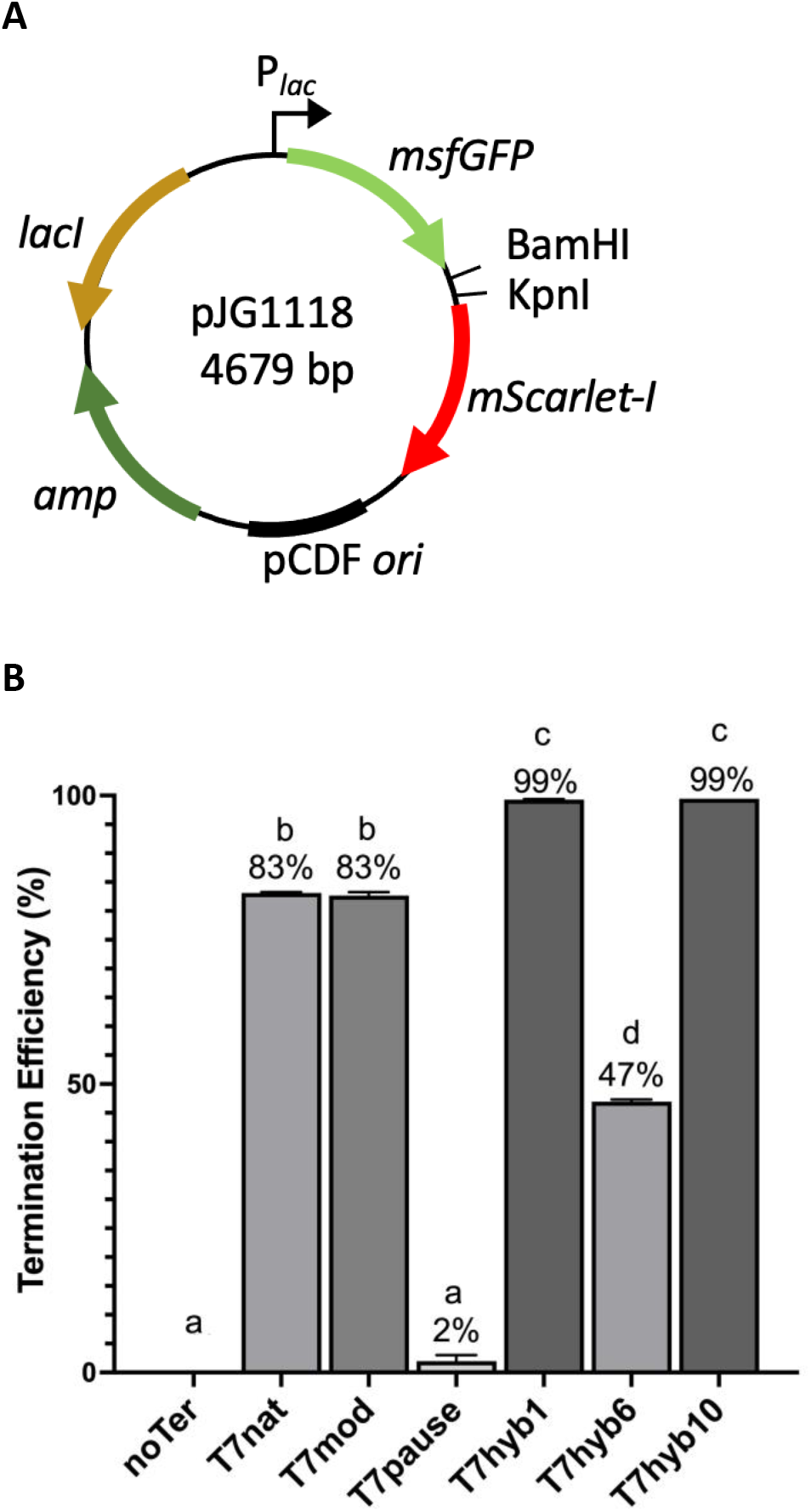
In vivo performance of selected terminators with the bacterial polymerase EcRNAP. (A) Terminator sequences were placed between fluorescent protein-encoding genes located downstream of the *lac* promoter, using *Bam*HI and *Kpn*I restriction sites. Plasmids were transformed into *E. coli* MG1655 and reporter gene expression was used to assess terminator efficiency. (B) Bar chart of the mean termination efficiency of engineered terminators for EcRNAP. Error bars represent standard deviation from the mean. Different letters denote statistically significant differences (P < 0.05) according to a Tukey multiple comparison test.

## DATA AND REAGENT STATEMENT

Bacterial strains and plasmids are available upon request. The data and reagents underlying this article are available within the article and in the supplemental material section.

## ACKNOWLEDGEMENTS

Financial support for this work was provided by the National Institutes of Health (NIH) grant 1R15GM132852-01.

